# Conserved Cx_n_C motifs in Kaposi’s sarcoma-associated herpesvirus ORF66 are required for viral late gene expression and mediate its interaction with ORF34

**DOI:** 10.1101/728139

**Authors:** Allison L. Didychuk, Angelica F. Castañeda, Lola O. Kushnir, Carolyn J. Huang, Britt A. Glaunsinger

## Abstract

Late gene transcription in the beta- and gammaherpesviruses depends on a set of virally-encoded transcriptional activators (vTAs) that hijack the host transcriptional machinery and direct it to a subset of viral genes that are required for completion of the viral replication cycle and capsid assembly. In Kaposi’s sarcoma-associated herpesvirus (KSHV), these vTAs are encoded by ORF18, ORF24, ORF30, ORF31, ORF34, ORF66. Assembly of the vTAs into a complex is critical for late gene transcription, and thus deciphering the architecture of the complex is central to understanding its transcriptional regulatory activity. Here, we generated an ORF66-null virus and confirmed that it fails to produce late genes and infectious virions. We show that ORF66 is incorporated into the vTA complex primarily through its interaction with ORF34, which is mediated by a set of four conserved cysteine-rich motifs in the C-terminal domain of ORF66. While both ORF24 and ORF66 occupy the canonical K8.1 late gene promoter, their promoter occupancy requires the presence of the other vTAs, suggesting that sequence-specific, stable binding requires assembly of the entire complex on the promoter. Additionally, we find that ORF24 expression is impaired in the absence of a stable vTA complex. This work extends our knowledge about the architecture of the KSHV vPIC and suggests that it functions as a complex to recognize late gene promoters.

**IMPORTANCE:** Kaposi’s sarcoma-associated herpesvirus (KSHV; human herpesvirus 8) is an oncogenic gammaherpesvirus that is the causative agent of multiple human cancers. Release of infectious virions requires production of capsid proteins and other late genes, whose production are transcriptionally controlled by a complex of six virally-encoded proteins that hijack the host transcription machinery. It is poorly understood how this complex assembles or what function five of its six components play in transcription. Here, we demonstrate that ORF66 is an essential component of this complex in KSHV and that its inclusion in the complex is mediated through its C-terminal domain, which contains highly conserved cysteine-rich motifs reminiscent of zinc finger motifs. Additionally, we examine assembly of the viral pre-initiation complex at late gene promoters and find that while sequence-specific binding of late gene promoters requires ORF24, it additionally requires a fully assembled viral pre-initation complex.

## INTRODUCTION

Gammaherpesviruses such as Kaposi’s sarcoma-associated herpesvirus (KSHV) and Epstein Barr virus (EBV), along with betaherpesviruses such as human cytomegalovirus (HCMV), are dsDNA viruses that co-opt and exploit endogenous cellular transcription machinery to facilitate viral gene expression. During the lytic phase of the lifecycle, gammaherpesviral genes are expressed in a temporal cascade starting with immediate early genes, followed by early, then late genes. In all classes of viral genes, transcription depends upon host cellular RNA polymerase II (Pol II). Immediate early and early genes possess promoters similar to host promoters, with canonical TATA boxes. In contrast, late genes have minimal promoters, characterized by the presence of a non-canonical TAT<u>T</u> box, which in KSHV is followed by an RVNYS motif (1-3).

Late genes in beta- and gammaherpesviruses are transcribed by a set of virally encoded genes termed the viral transcriptional activators (vTAs), which form a complex termed the viral pre-initiation complex (vPIC) (4-6). Homologs of this complex are absent in alphaherpesviruses, which are thought to control late gene transcription by a distinct mechanism that depends on a canonical TATA box and Inr element (7). In KSHV, the vTAs are encoded by ORF18, ORF24, ORF30, ORF31, ORF34, and ORF66. Stop mutants of five of the six KSHV vTAs have been generated and tested, revealing that they share a common phenotype in which late gene transcription fails to occur, ultimately preventing release of infectious virions (6, 8-10). Disrupting protein-protein contacts within the complex also prevents late gene transcription, as preventing the incorporation of even the smallest vPIC component, ORF30 (77 amino acids) by disrupting its interaction with its binding partner ORF18 completely prevents late gene transcription in both KSHV and HCMV (11, 12). Although deletion of KSHV ORF66 has not been tested, an EBV mutant lacking the ORF66 homolog (BFRF2) fails to produce infectious virions due to a defect in late gene transcription (4). Thus, the six viral components of the vPIC are absolutely required for late gene transcription.

It is well-established that the six vTAs form a complex. One of these vTAs, ORF24, binds late gene promoter DNA and directly recruits Pol II to facilitate late gene expression (6). However, the function of the vTA complex as a whole beyond polymerase recruitment is unknown. ORF24 and its homologs are considered to be viral TATA-binding proteins (vTBP), as they have weak sequence similarity to host TBP and *in silico* modeling suggests they may structurally mimic TBP (13). However, there is a dearth of structural or functional information for the remaining five vTAs.

Here, we confirm that KSHV ORF66 is essential for infectious virion production due to its role in late gene transcription. We demonstrate that ORF66 interacts with ORF18, ORF31, and ORF34 and that its interaction with ORF34, but not ORFs 18 and 31, is mediated by the C-terminal domain of ORF66. Disruption of conserved cysteine-rich motifs within the C-terminal domain of ORF66 prevents late gene transcription due to disruption of the interaction between ORF66 and ORF34. We also demonstrate that stable binding of ORF24 on late gene promoters requires both ORF66 and ORF30. These results extend our understanding of the architecture of the vTA complex as well as provide novel insights into its ability to recognize and bind late gene promoters *in vivo*.

## RESULTS

### ORF66 is essential for late gene expression in KSHV

KSHV ORF66 is a conserved protein with homologs in all beta and gammaherpesviruses. Based on the phenotype observed upon deletion or mutation of the other KSHV vPIC components (6, 8-10), we predicted that ORF66 would similarly be essential for viral replication and late gene expression in KSHV. We generated an ORF66-deficient recombinant KSHV BAC16-derived virus (ORF66.stop) using the Red recombinase system (14). As the coding region of ORF66 partially overlaps with that of ORF67, we inserted two adjacent stop codons at amino acids 25 and 26 of ORF66 (**Figure 1A**). We also engineered a corresponding mutant rescue (ORF66.MR) to ensure that any phenotypes observed were not due to secondary mutations elsewhere in the BAC. The sequence of the recombinant BACs was confirmed by Sanger sequencing, and they were digested with *RsrII* and *SbfI* to ensure that no large-scale recombination had occurred during mutagenesis (**Figure 1A-B**). We then generated latently infected iSLK.BAC16 cell lines by transfecting BACs into HEK293T cells followed by co-culture with SLK-puro cells harboring a doxycycline-inducible copy of ORF50 (RTA) to allow for efficient reactivation.

**Figure 1.**
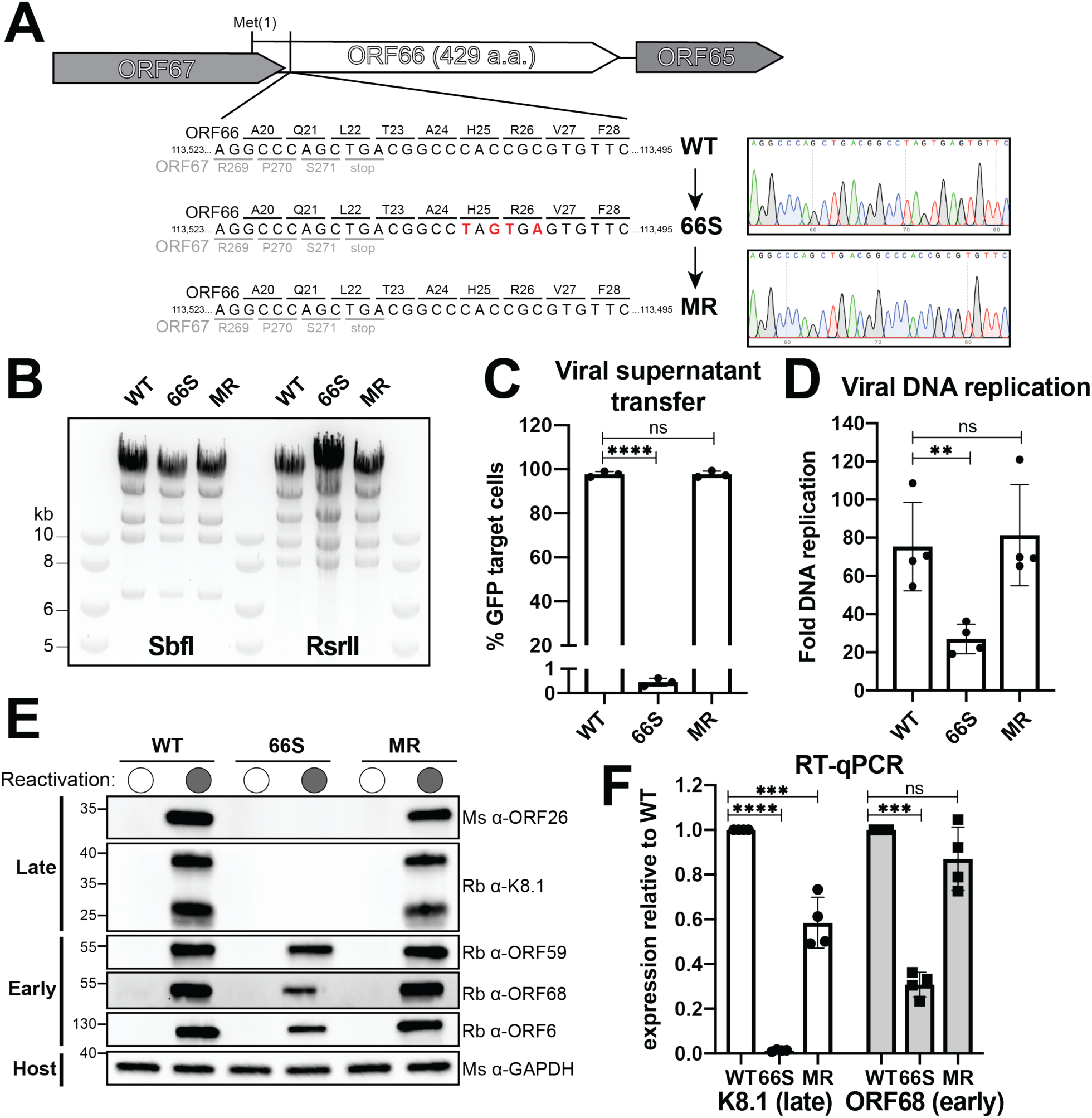
ORF66 is essential in KSHV and required for late gene transcription. A) Diagram showing the genomic locus of ORF66 with surrounding genes ORF67 (which partially overlaps ORF66) and ORF65, depicting the location of introduced mutations. Mutations were confirmed by Sanger sequencing (right). B) Digestion of the recombinant BACs with SbfI or RsrII demonstrates that introduction of mutations did not introduce large-scale changes. C) Infectious virion production was measured by supernatant transfer from reactivated iSLK cell lines followed by flow cytometry. Data are from three independent biological replicates with statistics calculated using an unpaired t-test, where (****) p < 0.0001. D) Viral DNA replication was measured using qPCR before and after reactivation. Data are from four independent biological replicates with statistics calculated using an unpaired t-test, where (**) p < 0.01. E) Western blotting of whole cell lysate (20 μg) reveals that early genes are largely unaffected by the ORF66.stop mutation, but late gene products cannot be detected. F) RT-qPCR reveals that the defect observed in (E) are due to a transcriptional effect caused by the absence of ORF66. Data are from four independent biological replicates with statistics calculated using an unpaired t-test, where (****) p < 0.0001, (***) p < 0.001, and (**) p < 0.01.

To assess whether ORF66 is essential for the completion of the viral life cycle, we monitored production of infectious progeny virions using a supernatant transfer assay. The BAC16 system contains a constitutively expressed GFP reporter gene, allowing for quantitation of infected target cells using flow cytometry. No detectable virus was produced by the ORF66.stop cell line, whereas the WT and ORF66.MR cell lines produced sufficient virus to infect nearly all of the target cells (**Figure 1C**). Thus, ORF66 is required for completion of the viral replication cycle.

We next tested whether ORF66 plays a role in replication of the viral genome. It is well established that late gene transcription is licensed by the initiation of viral genome replication (15-17). Although it has been previously reported that other vTA mutants do not exhibit a defect in viral genome replication (6, 8-10), we recently reported that mutations in ORF24 result in a ∼6-fold defect in viral genome replication (1). Similarly, we find that the ORF66.stop virus has a modest ∼3-fold defect in viral genome replication that is rescued in the ORF66.MR virus (**Figure 1D**).

To further evaluate the role(s) of ORF66 in the viral replication cycle, we examined expression of representative KSHV early and late genes in lytically reactivated iSLK cells containing WT, ORF66.stop, or ORF66.MR viruses by Western blot (**Figure 1E**) and RT-qPCR (**Figure 1F**). While the ORF66.stop infection produced the early proteins ORF59, ORF6, and ORF68, the late proteins K8.1 and ORF26 were not detectable (**Figure 1E**). In contrast, both early and late proteins were expressed in the WT and ORF66.MR infected cells (**Figure 1E**). RT-qPCR-based measurements of viral RNA from the ORF68 (early) and K8.1 (late) loci confirmed that the selective absence of late proteins in the ORF66.stop infections was due to a transcriptional defect (**Figure 1F**). The moderate decrease in both the transcript and protein of early gene ORF68 is consistent with the observation that most viral transcripts are downregulated in the absence of the vPIC component ORF24 (1). We also observed less ORF6 (the ssDNA binding protein involved in viral DNA replication) in the ORF66.stop samples, which could explain the reduced levels of viral DNA replication observed in the ORF66.stop cell line (**Figure 1D-E**). Thus, while ORF66 modestly contributes to KSHV early gene expression and DNA replication, it is essential for late gene expression.

### ORF66 is a component of the KSHV vTA complex

Based on our observation that ORF66 is essential for late gene transcription and our previous observation that ORF66 interacts with ORF18 (11), we sought to further characterize the interactions of ORF66 within the vTA complex (**Figure 2A**). We began by assessing its association with the complex as a whole upon immunoprecipitation of different vTA components in transiently transfected HEK293T cells. We used FLAG magnetic beads to enrich for a FLAG-tagged ORF and tested whether the remaining five Strep-tagged vTA complex components could be co-immunoprecipitated (**Figure 2B**). As we previously showed, the vTA complex can be isolated by immunoprecipitation of ORF18 (**Figure 2B**). We also tested if the complex could be isolated by immunoprecipitation of ORF31 or ORF66. Notably, although ORF31 was expressed as well as ORF18, it co-immunoprecipitated less ORF18 (and thus, less ORF30) relative to immunoprecipitation by ORF18 or ORF66 (**Figure 2B**). Overall, however, the structural integrity of the vTA complex is highlighted by the observation that immunoprecipitation of either ORF18, ORF31, or ORF66 resulted in co-purification of the remaining five vTA components.

**Figure 2.**
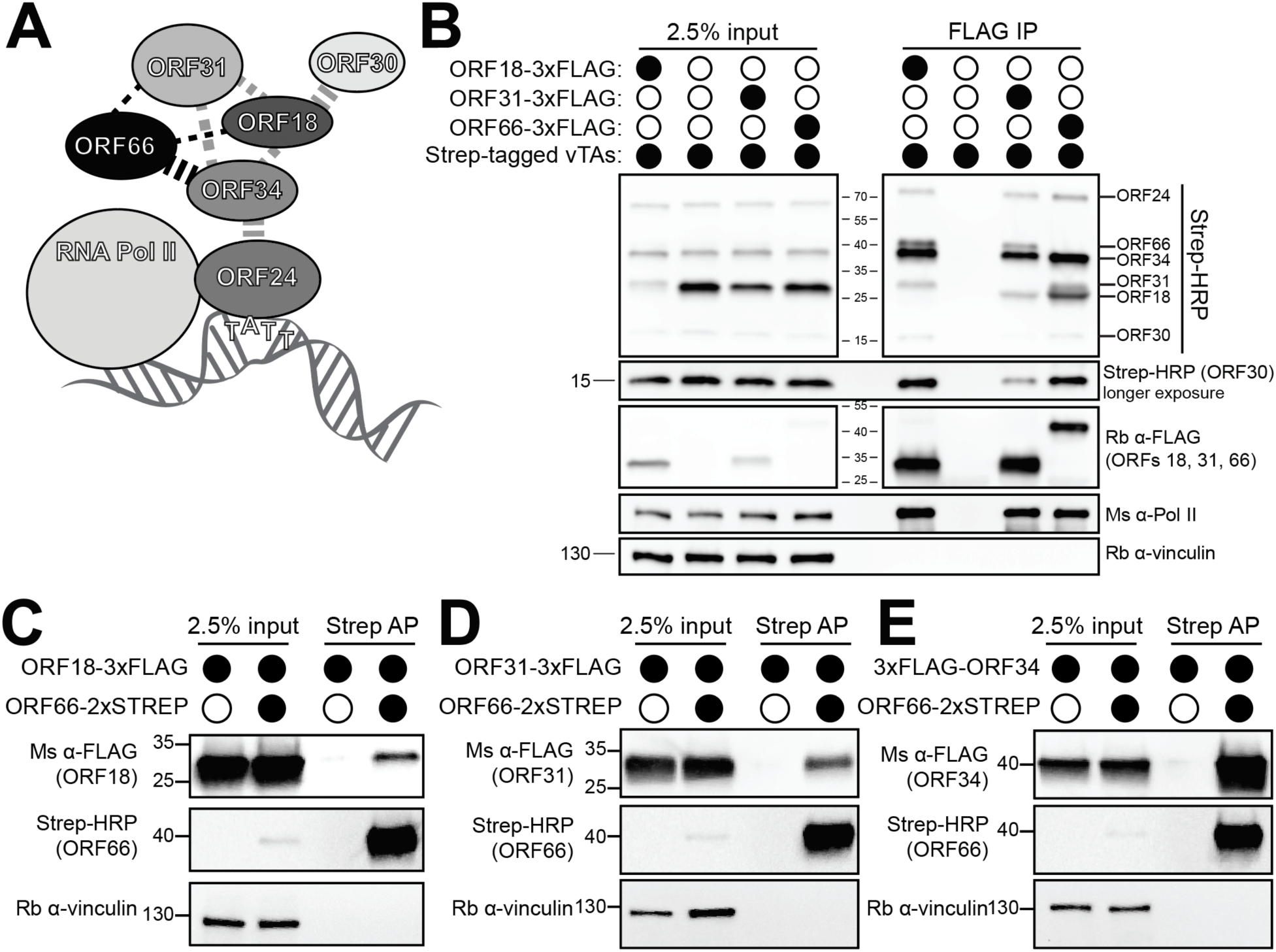
ORF66 is a component of the vPIC complex and directly interacts with ORFs 18, 31, and 34. A) Diagram of the vPIC components in KSHV. Line weights indicate relative strength of interactions. B) ORF18, ORF31, and ORF66 can immunoprecipitate the entire vPIC complex. HEK293T cells were transiently transfected with FLAG- or Strep-tagged vTAs and a co-IP was performed using anti-FLAG magnetic beads. In (C-E), StrepTactinXT magnetic beads were used to isolate Strep-tagged ORF66 and demonstrate ORF18 (C), ORF31 (D), and ORF34 (E) co-IP with ORF66.

To assess the contacts that ORF66 makes within the vTA complex, we tested its pairwise interactions with other components by co-IP. As has been previously observed, ORF66 interacts with ORF18 (**Figure 2C**) (11), ORF31 (**Figure 2D**), and most robustly with ORF34 (**Figure 2E**) (10). We were unable to detect direct protein-protein interactions between ORF66 and ORF24 or ORF30 (data not shown). Thus, like ORFs 18 and 34, ORF66 exhibits multiple interactions within the vTA complex.

### The interaction between ORF66 and ORF34 is mediated by the C-terminal domain of ORF66

To define the region(s) of ORF66 responsible for mediating its vTA protein-protein interactions, we first created truncations of ORF66 that roughly divide the protein into two domains, and then tested each for interaction with its binding partners by co-immunoprecipitation (**Figure 3A**). Interestingly, both the N- and C-terminal domains of ORF66 can still interact to varying degrees with ORF18 (**Figure 3B**) and ORF31 (**Figure 3C**). In contrast, the interaction with ORF34 mapped exclusively to the ORF66 C-terminal domain (**Figure 3D**). The robust interaction between ORF34 and ORF66 (in comparison to the significantly weaker interactions between ORF66 and ORF18 or ORF31) suggests that this interaction may drive incorporation of ORF66 into the vTA complex.

**Figure 3.**
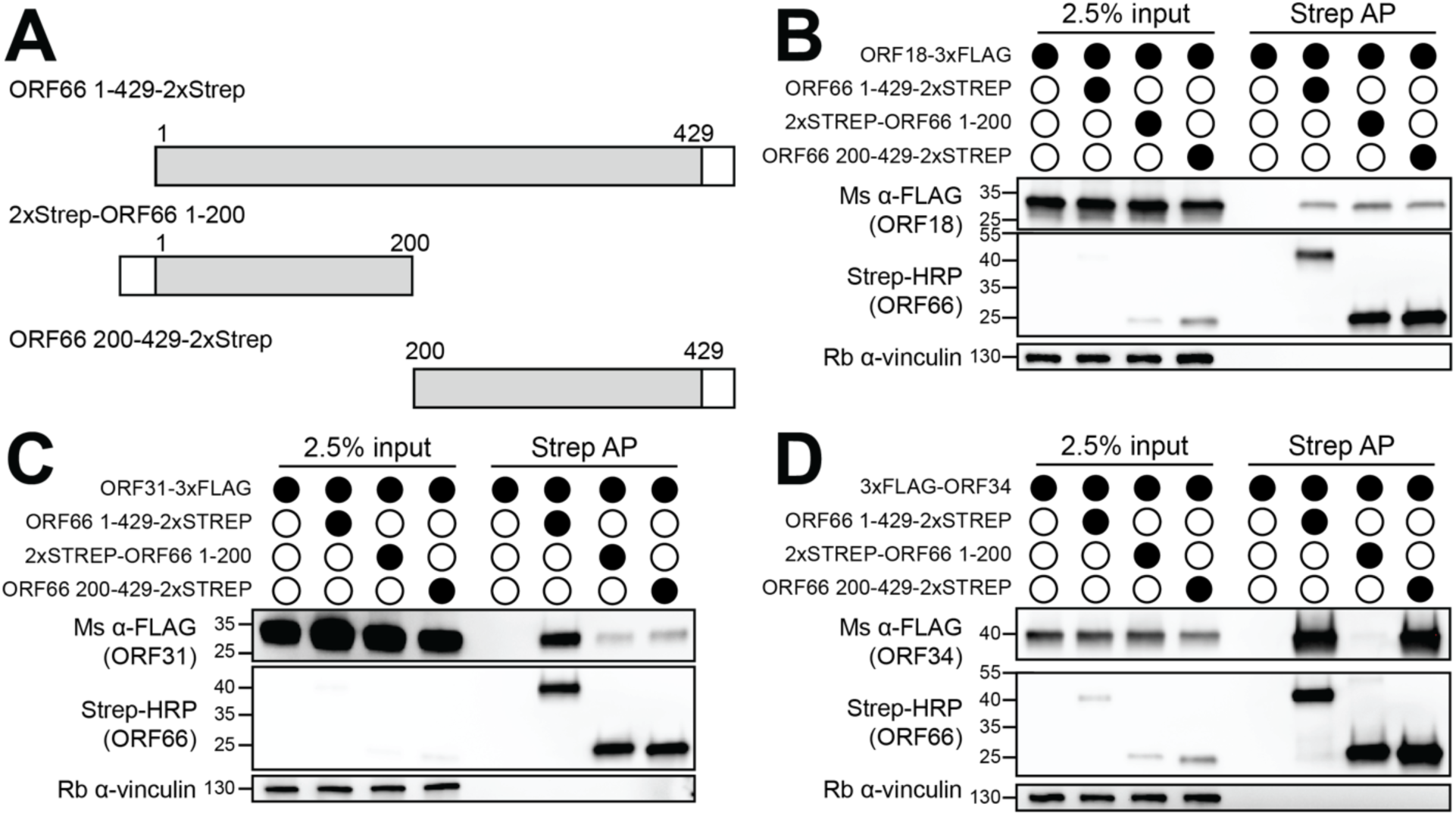
The C-terminal domain of ORF66 mediates protein-protein interactions within the vPIC. A) Diagram showing constructs used to test the domain structure of ORF66. B-D) HEK293T cells were transiently transfected with Strep-tagged ORF66 and the indicated FLAG-tagged vTA, then co-immunoprecipitated with StrepTactinXT beads (Strep AP) followed by Western blotting.

### The C-terminal domain of ORF66 contains conserved Cx_n_C motifs required for late gene expression

We next sought to further refine which residues underlie the robust interaction between the C-terminal domain of ORF66 and ORF34. We performed a multiple sequence alignment of the C-terminal domain of KSHV ORF66 and its homologs in the gammaherpesviruses MHV68 (mu66) and EBV (BFRF2) along with homologs from the betaherpesviruses MCMV (M49), HCMV (UL49), HHV6A (U33), and HHV7 (UL49) using T-coffee (18) (**Figure 4A**). It was immediately striking that four Cx_n_C motifs (motifs I-IV) were perfectly conserved in the aligned region in all available beta and gammaherpesvirus homologs of ORF66.

**Figure 4.**
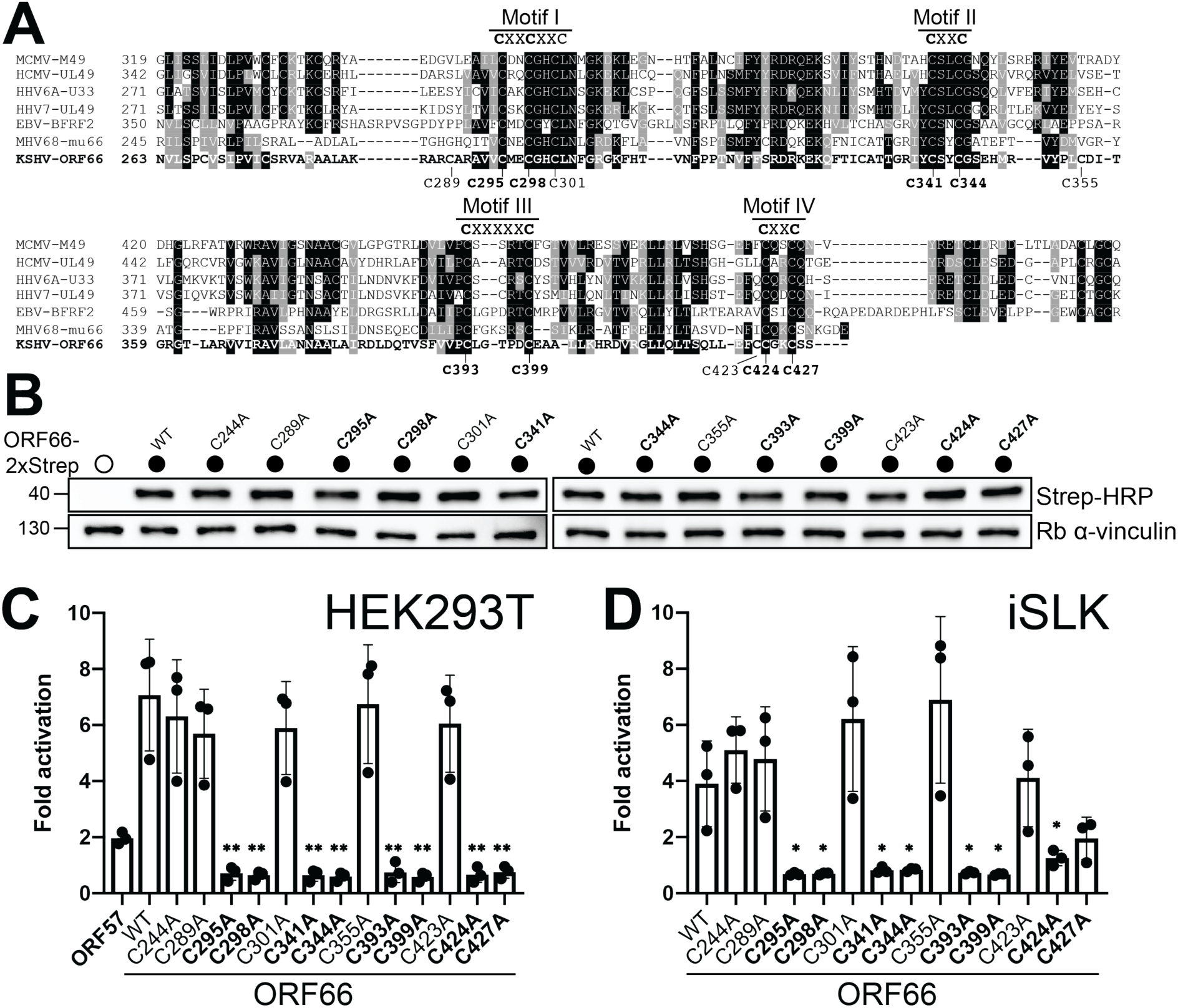
ORF66 contains conserved Cx_n_C motifs essential for late gene transcription. A) Multiple sequence alignment between KSHV ORF66 and homologs from related beta- and gammaherpesviruses. Cysteine residues used in subsequent experiments are indicated below the alignment, and presence of conserved CxnC motifs is indicated above the alignment. Bold indicates cysteines that affect late gene transcription. B) Western blot of whole cell lysate (15 μg) from HEK293T cells transiently transfected with plasmids containing wild-type or mutant Strep-tagged ORF66. C) HEK293T cells were transfected with plasmids encoding the six vTAs (including either wild-type or mutant ORF66), the pGL4.16 firefly luciferase plasmid under control of either the K8.1 or ORF57 promoter, and the pRL-TK renilla luciferase plasmid as a transfection control. After 24 h, cell lysates were harvested and luciferase activity was measured. D) iSLK cells were transfected with plasmids encoding wild-type or mutant ORF66, the pGL4.16 luciferase plasmid under control of the K8.1 promoter, and the pRL-TK renilla luciferase plasmid control. In both (C) and (D), fold activation was normalized to a control with empty vector replacing the vTAs (C) or ORF66 (D). Data are from three independent biological replicates with statistics calculated using an unpaired t-test, where (**) p < 0.01 and (*) p < 0.05.

We individually mutated to alanine the nine conserved cysteines within motifs I-IV, along with four nonconserved cysteine residues interspersed between the conserved motifs (**Figure 4A**). Importantly, all of the ORF66 mutants were expressed comparably to the wild-type protein when transfected into HEK293T cells, suggesting that these residues, and hence the Cx_n_C motifs, are not critical for protein stability (**Figure 4B**).

To determine if the Cx_n_C motifs were necessary for late gene transcription, we used a previously described reporter assay wherein firefly luciferase gene is under the control of an early gene promoter (from ORF57) or a late gene promoter (K8.1) (11). In addition to the firefly luciferase reporter, the six vTAs (including wild-type or mutant ORF66) were transiently transfected into HEK293T cells along with a renilla luciferase reporter to control for transfection efficiency. Using this assay, we found that eight of the nine conserved cysteines are required for late gene transcription (**Figure 4C**). Although cysteine 301 (C301) is perfectly conserved and is part of the extended CxxCxxC in motif I, its mutation does not affect late gene transcription (**Figure 4C**).

We then tested the role of these ORF66 cysteine residues in the context of KSHV infection using a modified version of the assay described above. The iSLK ORF66.stop cell line was reactivated from latency then transfected with the luciferase reporter plasmids along with a plasmid containing wild-type or mutant ORF66. In this assay, the remaining components of the vPIC are supplied by the viral genome. The results in infected cells paralleled those obtained in 293T cells, as the same eight conserved cysteine residue mutants in motifs I-IV (C295A, C298A, C341A, C344A, C393A, C399A, C424A, C427A) selectively failed to activate the luciferase late gene reporter (**Figure 4D**). In contrast, C301A in motif I and the four cysteine mutants interspersed between motifs I-IV activated the late promoter similar to wild-type ORF66 (**Figure 4D**). In conclusion, our results show that with the exception of C301 at the end of motif I, the conserved cysteines comprising the four Cx_n_C motifs in the C-terminal domain of ORF66 are critical for late gene transcription.

### The Cx_n_C motifs in the C-terminal domain of ORF66 are required for interaction with ORF34 and its incorporation into the vPIC

We next evaluated what role the Cx_n_C motifs in ORF66 might play in assembly of the vPIC. We selected representative cysteine mutants from the four motifs (C295, C341, C393, C424) and tested their ability to interact with each of the ORF66 vPIC binding partners (**Figure 5A-C**). All of the mutants retained wild-type levels of binding to ORF18 or ORF31 (**Figure 5A-B**). In contrast, motif I (C295A) and II (C341A) mutants displayed greatly reduced binding to ORF34, and motif III (C393A) and IV (C424A) mutants failed to interact with ORF34 (**Figure 5C**). This is consistent with our previous observation that the C-terminal domain of ORF66 mediates the interaction with ORF34.

**Figure 5.**
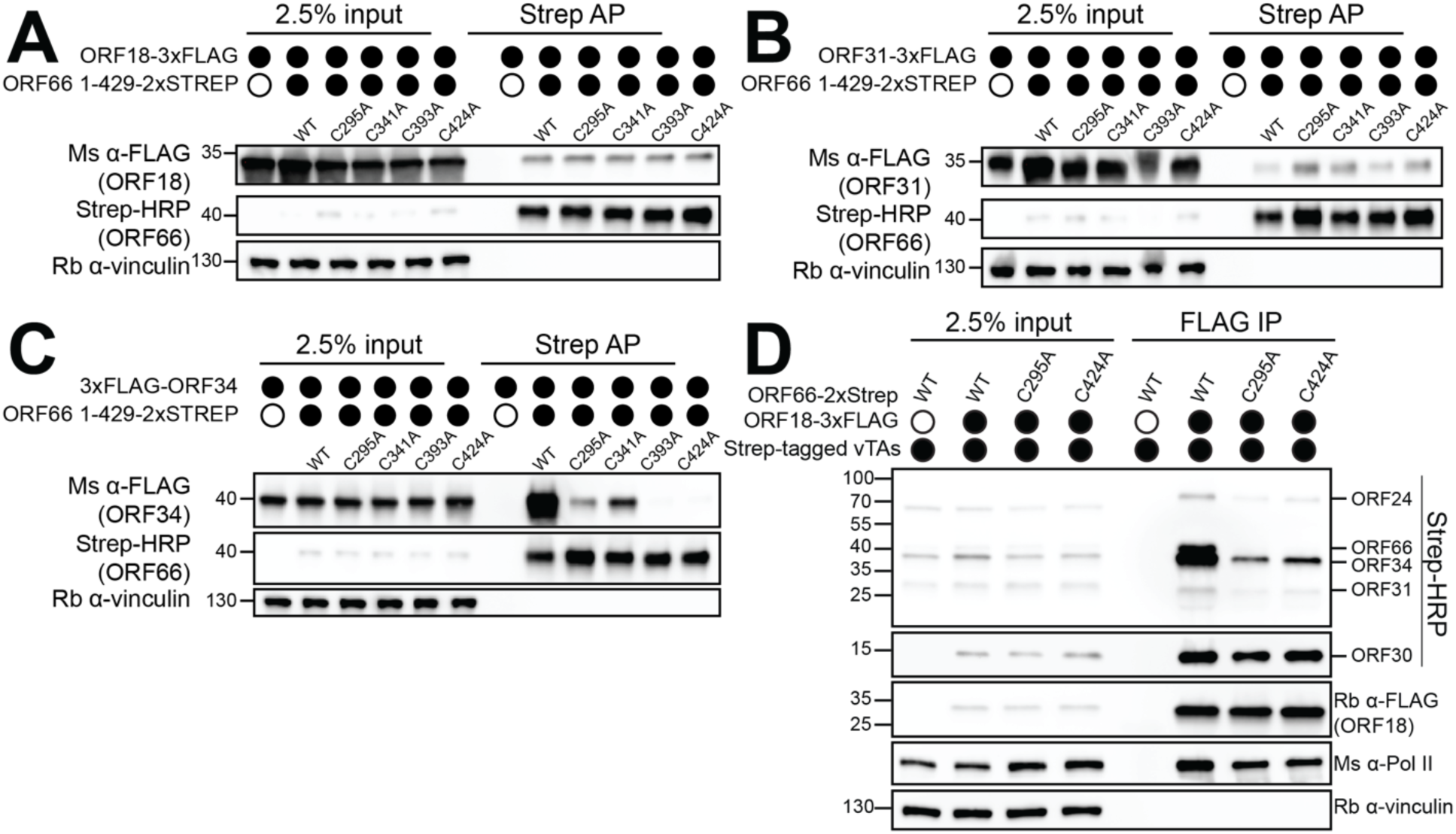
The Cx_n_C motifs in ORF66 are required for interaction with ORF34. A-C) HEK293T cells were transiently transfected with wild-type or mutant Strep-tagged ORF66 and the indicated FLAG-tagged vTA, then co-immunoprecipitated with StrepTactinXT beads followed by Western blotting. D) HEK293T cells were transiently transfected with wild-type or mutant Strep-tagged ORF66 along with FLAG-tagged ORF18, ORF24, ORF30, ORF31, and ORF34. A co-IP with anti-FLAG magnetic beads was performed followed by Western blotting.

We then tested if ORF66 mutants with significantly weakened (i.e. motif I C295A) or fully impaired (i.e. motif IV C424A) ORF34 binding impacted incorporation of ORF66 into the vTA complex (**Figure 5D**). We isolated the vTA complex by immunoprecipitation of Flag-tagged ORF18 in the presence of the remaining 5 strep-tagged vTA proteins, including wild-type or mutant ORF66. Neither ORF66 C295A nor C424A were incorporated into the vTA complex (**Figure 5D**), despite the fact that these mutants retained pairwise interactions with ORF18 and ORF31 (see Figure 2). Notably, the remaining vTA components (ORFs 18, 24, 30, 31, and 34) still assembled into the complex in the absence of ORF66. Thus, the robust ORF66 Cx_n_C motif-mediated interaction with ORF34, but not the weaker interactions of ORF66 with ORF18 and ORF31, is essential to recruit ORF66 into the vTA complex.

### An intact vPIC is required for stable binding of ORF66 and ORF24 at late gene promoters *in vivo*

ORF24 is the only vTA known to directly contact promoter DNA, although other vTAs likely co-localize at late gene promoters via protein-protein interactions within the complex (1, 6). However, given that ORF66 possesses Cx_n_C motifs, which are frequently found in nucleic acid binding proteins (19), we considered that it might independently bind promoter DNA. We also sought to test whether other vTAs, including ORF66, contribute to ORF24 promoter specificity or binding during infection.

In this regard, we generated KSHV containing C-terminally HA tagged ORF66 in an otherwise WT BAC16 background (66HA) or in a BAC16 lacking ORF24 (66HA/24S) to evaluate whether ORF66 associates with the late gene promoter and, if so, whether its association requires ORF24. We also generated KSHV with N-terminally HA tagged ORF24 (HA24; (1)) and lacking either ORF66 (HA24/66S) or ORF30 (H24/30S) to determine whether promoter binding by ORF24 is influenced by other vTAs. Unlike ORF66, the ORF30 vTA is a small protein with no predicted nucleic acid binding properties and its only connection with the complex occurs through ORF18 (see **Figure 2A**) (11). Thus, ORF30 deletion should enable evaluation of the general importance of the vTA complex in ORF24 late gene promoter binding. We generated the recombinant BACs and iSLK cell lines as described earlier and digested the BACs with *RsrII* and *SbfII* to ensure that no large-scale recombination occurred during mutagenesis (**Figure 6A**).

**Figure 6.**
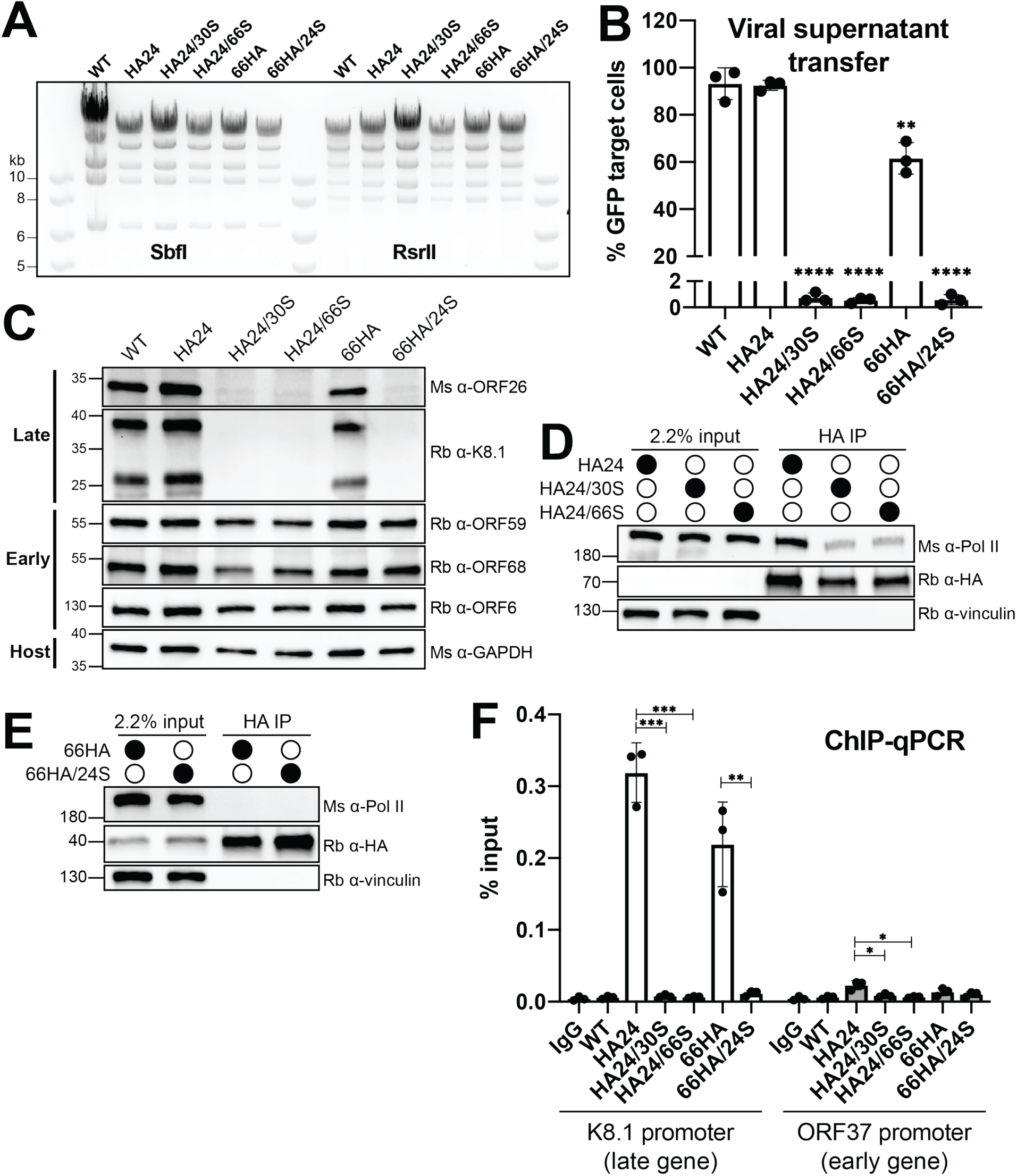
ORF24 does not bind to late gene promoters in the absence of ORF30 or ORF66. iSLK cell lines were created using the recombinant BAC16 system. HA tags were added to the endogenous copies of the N-terminus of ORF24 (HA24) or the C-terminus of ORF66 (66HA). In select BACs, ORF24, ORF30, or ORF66 were deleted by the introduction of a stop codon early in the ORF (24S, 30S, and 66S respectively). A) Digestion of the recombinant BACs with SbfI or RsrII demonstrates that recombination did not introduce large-scale changes. B) Infectious virion production was measured by supernatant transfer from reactivated iSLK cell lines followed by flow cytometry. Data are from three independent biological replicates with statistics calculated using an unpaired t-test, where (****) p < 0.0001 and (**) p < 0.01. C) Western blot of whole cell lysate (25 μg) from iSLK cell lines showing the relative levels of representative early and late genes. D) Western blots showing expression of HA-ORF24 in different cell lines. Proteins were immunoprecipitated with HA beads to enrich for ORF24, which is low abundance. E) Western blots showing expression of ORF66-HA in different cell lines. Proteins were immunoprecipitated with HA beads to enrich for ORF66, which is low abundance. F) ChIP-qPCR from the indicated cell lines was performed using an anti-HA antibody. The associated DNA from either the K8.1 promoter (a late gene promoter) or the ORF37 promoter (an early gene promoter) was quantified using promoter-specific primers. Data are from three independent biological replicates with statistics calculated using an unpaired t-test, where (***) p < 0.001, (**) p < 0.01, and (*) p < 0.05.

We first characterized each of the infected cell lines for their ability to produce infectious virions and express early and late genes upon lytic reactivation. As expected, iSLK cells containing HA24 KSHV and 66HA KSHV produced infectious virus as measured using a supernatant transfer assay, although in the case of 66HA the levels were modestly reduced (by ∼40%) relative to WT KSHV (**Figure 6B**). This reduction could be caused by an effect of the tag on ORF66 stability or by the location of the epitope tag in the viral genome, as it changes the 5’UTR of neighboring gene ORF65. In contrast, KSHV mutants lacking the individual vTAs (HA24/30S, HA24/66S, or 66A/24S) produced no detectable infectious virions (**Figure 6B**).

Western blotting confirmed that all of the engineered viruses expressed the representative early proteins ORF59, ORF68, and ORF6 (**Figure 6C**). However, the late proteins ORF26 and K8.1 were not produced in cell lines lacking any of the vTAs (**Figure 6C**). Late proteins were produced in the 66HA iSLK cells, although in agreement with the virion production data, their levels were slightly reduced relative to KSHV containing untagged ORF66 (WT) (**Figure 6C**). Finally, we confirmed expression of the HA-tagged ORF24 and ORF66 by immunoprecipitation with anti-HA beads (**Figure 6D-E**), which was necessary as both proteins are low abundance and cannot be easily detected in whole cell lysate. Interestingly, the levels of HA-ORF24 (but not ORF66-HA) were reduced upon deletion of either ORF30 or ORF66, suggesting that ORF24 expression is bolstered by an intact vTA complex (**Figure 6D**).

We next performed chromatin immunoprecipitation (ChIP) using the HA tag on endogenous ORF24 or ORF66 and quantified the amount of associated DNA by qPCR. As anticipated, both HA-ORF24 and ORF66-HA bound to the promoter of the K8.1 late gene but not of the early gene ORF37 (**Figure 6F**). Notably, ORF66-HA binding to the K8.1 promoter did not occur in the absence of ORF24 (66HA/24S), suggesting that it does not independently bind the late gene promoter (**Figure 6F**). Surprisingly, we detected no HA-ORF24 binding at the K8.1 promoter during infection with viruses lacking either ORF66 or ORF30 (**Figure 6F**). It is possible that the ChIP assay is not sensitive enough to detect DNA associated with the reduced levels of HA-ORF24 in these cells (**Figure 6D**). Alternatively, although ORF24 alone displays sequence specific DNA binding *in vitro* (6), its stable association with late promoters in cells may require an intact vTA complex. In summary, our results suggest that sequence-specific binding of ORF24 at late gene promoters is bolstered by the vTA complex.

## DISCUSSION

Here, we demonstrate that KSHV ORF66, similar to the other 5 vTAs (6, 8-10), is necessary for completion of the lytic replication cycle due to its critical role in late gene transcription. Additionally, our results complement previous work **(11)** in demonstrating that disruption of any of the protein-protein interactions within the complex, even when the remaining contacts between other vTA complex components are maintained, prevents late gene transcription. Despite recent progress in understanding the overall organization of the vTA complex, the role of each vTA is poorly understood. We mapped the interactions between ORF66 and other members of the vTA complex and identified four conserved Cx_n_C motifs within the C-terminal domain of ORF66. When mutated, these motifs abolish late gene transcription due to a drastic reduction in the ability of ORF66 to bind ORF34. We show that ORF66 is present at late gene promoters during infection, but does not bind in the absence of the vTBP mimic ORF24. ORF24 binding at late gene promoters requires both ORF66 and ORF30, suggesting the presence of all six vTAs at late gene promoters is necessary for stable binding of the vPIC.

The vTA complex displays remarkable physical and functional interconnectivity. While ORF34 was originally proposed to be the scaffold upon which the other vTAs assemble, it is clear that both ORF66 and ORF18 directly engage in interactions with three and four vTAs respectively, suggesting that the complex is stabilized by numerous protein-protein interactions between multiple core components. Furthermore, all six members of the vTA complex must be present and individual protein-protein interactions must be maintained in order for late gene transcription to occur. Disruption of the interaction between ORF66 and ORF34 or the interaction between ORF30 and ORF18 (11) not only changes the composition of the vTA complex (by preventing inclusion of ORF66 or ORF30, respectively) but also destabilizes the complex, even though multiple contacts between other vTAs exist. Ultimately, determining the individual structures of vTAs and the architecture of the complex at molecular resolution will be key to interpreting the roles of known protein-protein contacts and deciphering their functions in late gene transcription.

Three of the six vTAs – ORF18, ORF34, and ORF66 – contain conserved cysteine/histidine motifs that in the context of zinc finger domains are frequently found in nucleic acid binding proteins. However, the observation that ORF66 only associates with the K8.1 promoter in the presence of ORF24 suggests that it does not bind promoter DNA (at least not alone) and is more likely to be recruited indirectly through contacts with the vTA complex. Thus, although the Cx_n_C motifs in ORF66 are required for late gene expression, they appear to mediate an interaction with ORF34 rather than contributing to promoter-specific binding by the vPIC. Mutations within the Cx_n_C motifs did not change the expression or stability of ORF66 in cells or its interactions with ORF18 or ORF31, making it unlikely that these mutations result in global unfolding. Instead, we hypothesize that they serve to structurally stabilize a region of the protein that directly interacts with ORF34.

Although ORF66 expression was not dependent on the other vTAs, we found that ORF24 protein levels decreased in the absence of ORF66 or ORF30. Uncovering the links between ORF24 expression and the vTA complex could lead to new insights regarding the regulation of late gene transcription. For example, the vTA complex or one of its components could be responsible for stabilization of the low abundance ORF24 protein. Alternatively, a component(s) of the vTA complex could be important for correct localization of ORF24 to the nucleus, as has been observed for the KSHV DNA replication complex (20). Regardless of the mechanism involved, it is important to keep in mind that a reduction in ORF24 levels might contribute to defects observed in cell lines where the other vTAs are deleted (6, 8-10).

ORF24 has weak sequence similarity but predicted structural similarity to host TATA-binding protein (TBP) (13), although the degree to which it structurally and functionally mimics TBP is unknown. ORF24 binds a TATT-containing probe *in vitro* yet does not bind the probe when the TATT motif is mutated to CCCC (6). Its homolog from EBV, BcRF1, binds both TATT- and TATA-containing motifs *in vitro* but does not bind an unrelated probe (21). These observations suggest that ORF24 has some inherent sequence specificity for T/A-rich elements, similar to TBP. However, many promoters that lack canonical TATA box are bound by TBP *in vivo* (22), in contrast to the specificity for TATT(T/A)AAA+RVNYS promoters that are bound by the vPIC (1). Thus, the mechanism by which ORF24 specifically occupies the minimalistic late gene promoters remains a central unanswered question in viral late gene biology. We detected no binding of ORF24 at late gene promoters of viruses lacking either ORF30 or ORF66. These results suggest that despite apparent sequence specificity of ORF24 *in vitro* (6), sequence-specific binding during infection requires the other components of the vTA complex. Neither ORF66 nor ORF30 directly interact with ORF24, and thus regulation of ORF24 binding is presumably orchestrated through the web of protein-protein interactions within the vTA complex.

Despite significant progress in defining the components of the vTA complex and the protein-protein interactions in which they participate, the functional contribution of each vTA within the vPIC remains largely enigmatic. By analogy to the similarity between ORF24 and TBP, the remaining five vTAs may be mimicking other core host general transcription factors. Alternatively, the vTAs may be responsible for recruitment of host GTFs. Either of these possibilities allows for a contribution to sequence-specific binding by the non-ORF24 vTAs, although such binding may only be observed in the context of the fully assembled late gene vPIC.

## MATERIALS AND METHODS

### Plasmids

All plasmids described below were generated using InFusion cloning (Clontech) unless indicated otherwise; all have been deposited in Addgene. ORF66 was subcloned into the BamHI and XhoI sites of pcDNA4/TO-2xStrep (C-terminal tag) to generate pcDNA4/TO-ORF66-2xStrep (Addgene plasmid #130953). ORF66 aa 1-200 was cloned into the NotI and XhoI sites of pcDNA4/TO-2xStrep (N-terminal tag) to generate pcDNA4/TO-2xStrep-ORF66 1-200 (Addgene plasmid #130954) and ORF66 aa 200-429 was cloned into the BamHI and XhoI sites of pcDNA4/TO-2xStrep (C-terminal tag) to generate pcDNA4/TO-ORF66 200-429-2xStrep (Addgene plasmid #130955). Point mutations in pcDNA4/TO-ORF66-2xStrep (Addgene plasmids #131109-131121) were generated using inverse PCR site-directed mutagenesis with Phusion DNA polymerase (New England Biolabs) with primers as listed in Table 1. PCR products from inverse PCR were DpnI treated, ligated using T4 PNK and T4 DNA ligase and transformed into *Escherichia coli* XL-1 Blue cells. ORF66-2xStrep, ORF30-2xStrep, and ORF24-3xFlag were subcloned into the AgeI and EcoRI sites of pLJM1 that had been modified to confer zeocin resistance (Addgene plasmids #130957-130959). Plasmid K8.1 Pr pGL4.16 (Addgene plasmid #120377) contains the minimal K8.1 promoter and ORF57 Pr pGL4.16 (Addgene plasmid #120378) contains a minimal ORF57 early gene promoter and have been described previously (11). Plasmid K8.1 Pr pGL4.16+Ori (Addgene plasmid #131038) contains the left origin of replication along with a 100 bp fragment of the K8.1 promoter and has been described previously (1). Plasmids pcDNA4/TO-ORF18-2xStrep (Addgene plasmid # 120372), pcDNA4/TO-ORF24-2xStrep (Addgene plasmid #129742), pcDNA4/TO-ORF30-2xStrep (Addgene plasmid #129743), pcDNA4/TO-ORF31-2xStrep (Addgene plasmid #129744), pcDNA4/TO-2xStrep-ORF34 (Addgene plasmid #120376) have been previously described (11). Plasmid pRL-TK (Promega) was kindly provided by Dr. Russell Vance. Lentiviral packaging plasmids psPAX2 (Addgene plasmid #12260) and pMD2.G (Addgene plasmid #12259) were gifts from Dr. Didier Trono.

**TABLE 1.**
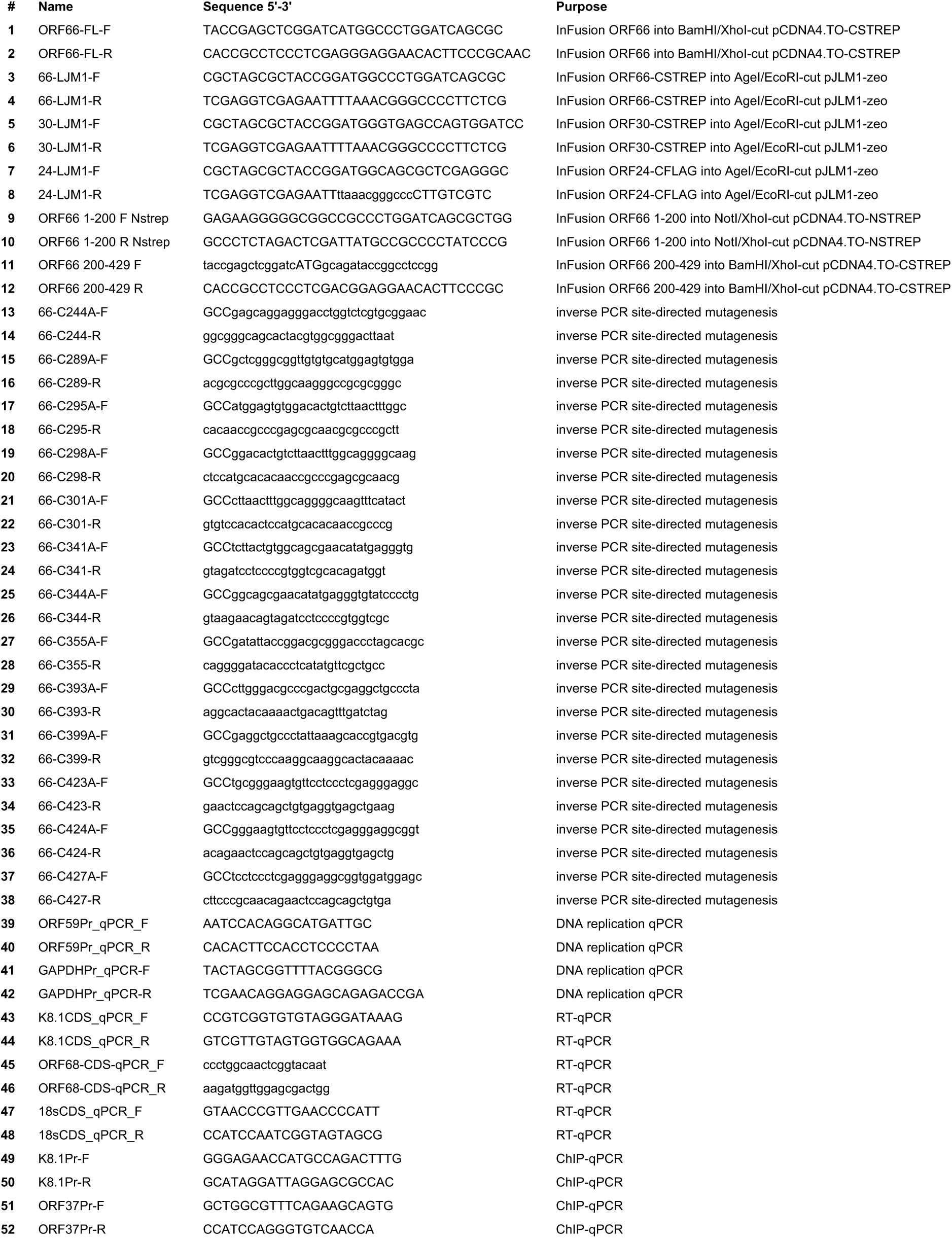

### Cell lines

HEK293T cells (ATCC CRL-3216) were maintained in DMEM supplemented with 10% FBS (Seradigm). HEK293T cells constitutively expressing ORF66-2xSTREP (HEK293T-ORF66), ORF24-3xFLAG (HEK293T-ORF24), ORF30-2xSTREP (HEK293T-ORF30), or 2xSTREP-ORF34 (HEK293T-ORF34) were maintained in DMEM supplemented with 10% FBS and 500 μg/ml zeocin.

iSLK-puro cells were maintained in DMEM supplemented with 10% FBS and 1 μg/ml puromycin. The iSLK cell line harboring the KSHV genome on the bacterial artificial chromosome BAC16 and a doxycycline-inducible copy of the KSHV lytic transactivator RTA (iSLK-BAC16) has been previously described (14). All iSLK BAC16 cell lines were maintained in DMEM supplemented with 10% FBS, 1 mg/mL hygromycin, and 1 μg/ml puromycin (iSLK-BAC16 media).

### Cell line establishment and viral mutagenesis

HEK293T cells stably expressing ORF66, ORF24, and ORF30 were generated by lentiviral transduction for the purpose of propagating KSHV deletion mutants lacking these essential genes. Lentivirus was generated in HEK293T cells by co-transfection of pLJM1-ORF66, -ORF24, or -ORF30 along with the packaging plasmids pMD2.G and psPAX2. After 48 h, the supernatant was harvested and syringe-filtered through a 0.45 μm filter (Millipore). The supernatant was diluted 1:2 with DMEM and polybrene was added to a final concentration of 8 μg/ml. 1 × 10^6^ freshly trypsinized HEK293T cells were spinoculated in a 6-well plate for 2 h at 1000 x g. After 24 h the cells were expanded to a 10 cm tissue culture plate and selected for 2 weeks in media supplemented with 500 μg/ml zeocin (Sigma).

All viral ORF mutants were generated using the scarless Red recombination system in BAC16 GS1783 *Escherichia coli* as previously described (14). The modified BACs were purified using a Nucleobond BAC 100 kit (Clontech). BAC quality was assessed by digestion with RsrII and SbfI (New England Biolabs). Latently infected iSLK cell lines with modified virus were generated by transfection of HEK293T cells (either WT or stably expressing the relevant essential viral ORF) with 5 μg BAC DNA using PolyJet (SignaGen). The following day, the transfected HEK293T cells were trypsinized and mixed 1:1 with freshly trypsinzed iSLK-puro cells and treated with 30 nM 12-O-tetradecanoylphorbyl-13-acetate (TPA) and 300 mM sodium butyrate for 4 days to induce lytic replication. iSLK cells were then selected in media containing 300 μg/mL hygromycin B, 1 μg/mL puromycin, and 250 μg/mL G418. The hygromycin B concentration was increased to 500 μg/mL and 1 mg/mL until all HEK293T cells died.

### Virus Characterization

For reactivation studies, 1 × 10^6^ iSLK cells were plated in 10 cm dishes for 16 h, then induced with 1 μg/ml doxycycline and 1 mM sodium butyrate for an additional 72 h. To determine the fold DNA induction in reactivated cells, the cells were scraped and triturated in the induced media, and 200 μL of the cell/supernatant suspension was treated overnight with 80 μg/ml proteinase K (Promega) in 1x proteinase K digestion buffer (10 mM Tris-HCl pH 7.4, 100 mM NaCl, 1 mM EDTA, 0.5% SDS) after which DNA was extracted using a Quick-DNA Miniprep kit (Zymo). Viral DNA fold induction was quantified by qPCR using iTaq Universal SYBR Green Supermix (BioRad) on a QuantStudio3 Real-Time PCR machine with primers for the KSHV ORF59 promoter and normalized to the level of GAPDH promoter (Table 1).

Infectious virion production was determined by supernatant transfer assay. Supernatant from induced iSLK cells was syringe-filtered through a 0.45 μm filter, then 2 mL of the supernatant was spinoculated onto 1 × 10^6^ freshly trypsinized HEK293T cells for 2 h at 1000 x g. After 24 h, the media was aspirated, the cells were washed once with cold PBS and crosslinked in 4% PFA (Electron Microscopy Services) diluted in PBS. The cells were pelleted, resuspended in PBS, and 50,000 cells/sample were analyzed on a BD Accuri 6 flow cytometer. The data were analyzed using FlowJo (30).

Total RNA and protein were isolated from reactivated iSLK cells at 72 hours. RNA was isolated using a Direct-Zol RNA Miniprep Plus kit (Zymo). Purified RNA was treated with TURBO DNase (ThermoFisher) then cDNA was synthesized using AMV reverse transcriptase (Promega). The cDNA was used for qPCR analysis using iTaq Universal SYBR Green Supermix (BioRad) and signals for each ORF were normalized to 18s rRNA. Protein samples were resuspended in lysis buffer [150 mM NaCl, 50 mM Tris-HCl pH 7.4, 1 mM EDTA, 0.5% NP-40, and protease inhibitor (Roche)], rotated for 30 min at 4°C, clarified by centrifugation at 21,000 x g for 10 min, then 25 μg of lysate was used for SDS-PAGE and western blotting in TBST (Tris-buffered saline, 0.2% Tween 20) using rabbit anti-K8.1 (1:10,000), rabbit anti-ORF59 (1:10,000), rabbit anti-ORF6 (1:10,000), rabbit anti-ORF68 (1:5000), mouse anti-ORF26 (Novus, 1:500), and mouse anti-GAPDH (Abcam, 1:1000). Rabbit anti-ORF59, anti-K8.1 sera, and anti-ORF6 sera were produced by the Pocono Rabbit Farm and Laboratory by immunizing rabbits against full length MBP-ORF59, MBP-K8.1, or MBP-ORF6 [gifts from Denise Whitby (23)]. Rabbit anti-ORF68 was previously described (24).

### Immunoprecipitation and western blotting

For all DNA transfections, HEK293T cells were plated and transfected after 24 h at ∼70% confluency with PolyJet (SignaGen). Cell lysates were prepared 24 h after transfection by washing and pelleting cells in cold PBS followed by resuspension in lysis buffer and rotation at 4°C for 30 min. For isolation of endogenous HA-tagged proteins, 3.5 × 10^6^ iSLK cells (iSLK BAC16 cell lines ORF66-HA, ORF66-HA/ORF24.stop, HA-ORF24, HA-ORF24/ORF66.stop, and HA-ORF24/ORF30.stop) were reactivated for 48 h with 5 μg/ml doxycycline and 1 mM sodium butyrate. Lysates were clarified by centrifugation at 21,000 x g for 10 min, then 1 mg (for pairwise interaction IPs), 1.5 - 2 mg (for the entire late gene complex IPs), or 1.5 mg (for HA-tagged ORF66 and ORF24) of lysate was incubated with pre-washed MagStrep “type3” XT beads (IBA) (for pairwise interaction IPs), M2 anti-FLAG magnetic beads (Sigma) (for the entire late gene complex IPs), or anti-HA magnetic beads (Pierce) overnight in 150 mM NaCl, 50 mM Tris-HCl pH 7.4. The beads were washed 3x for 5 min each with IP wash buffer (150 mM NaCl, 50 mM Tris-HCl pH 7.4, 0.05% NP-40) and eluted with 2x Laemmli sample buffer (BioRad). Lysates and elutions were resolved by SDS-PAGE and western blotted in TBST using the following primary antibodies: Strep-HRP (Millipore, 1:2500), rabbit anti-FLAG (Sigma, 1:3000), mouse anti-FLAG (Sigma, 1:1000), rabbit anti-Vinculin (Abcam, 1:1000), mouse anti-Pol II CTD clone 8WG16 (Abcam, 1:1000), or rabbit anti-HA (Cell Signaling, 1:1000). Following incubation with primary antibodies, the membranes were washed with TBST and incubated with the appropriate secondary antibody. The secondary antibodies used were the following: goat anti-mouse-HRP (1:5000, Southern Biotech) or goat anti-rabbit-HRP (1:5000, Southern Biotech).

### Late Gene Reporter Assay

For assays in HEK293T cells, 1 × 10^6^ cells were plated in 6-well plates and after 24 h each well was transfected with 900 ng of DNA containing 125 ng each of pcDNA4/TO ORF18-2xStrep, ORF24-2xStrep, ORF30-2xStrep, ORF31-2xStrep, 2xStrep-ORF34, wild-type or mutant ORF66-2xStrep (or as a control, 750 ng of empty pcDNA4/TO-2xStrep plasmid), with either K8.1 Pr pGL4.16 or ORF57 Pr pGL4.16, along with 25 ng of pRL-TK as an internal transfection control. For assays in iSLK cells, 5 × 10^5^ iSLK-ORF66.stop cells were plated in 6-well plates and after 24 h each well was reactivated with 5 μg/ml doxycycline and 1 mM sodium butyrate, followed immediately by transfection with 500 ng wild-type or mutant pcDNA4/TO-ORF66-2xStrep (or as a control, 500 ng of empty pcDNA4/TO-2xStrep plasmid), 475 ng K8.1 Pr pGL4.16+Ori and 25 ng pRL-TK. After 24 h (for HEK293T assays) or 48 h (for iSLK assays), cells were rinsed twice with PBS, lysed by rocking for 15 min at room temperature in 500 μL of Passive Lysis Buffer (Promega), and clarified by centrifugation at 21,000 x g for 2 min. 20 μL of the clarified lysate was added in triplicate to a white chimney well microplate (Greiner bio-one) to measure luminescence on a Tecan M1000 microplate reader using a Dual Luciferase Assay Kit (Promega). The firefly luminescence was normalized to the internal Renilla luciferase control for each transfection. All samples were normalized to the corresponding control containing empty plasmid.

### Chromatin immunoprecipitation (ChIP)

ChIP was performed on 15-cm plates of iSLK cells (iSLK BAC16 cell lines WT, ORF66-HA, ORF66-HA/ORF24.stop, HA-ORF24, HA-ORF24/ORF66.stop, and HA-ORF24/ORF30.stop) reactivated for 48 h with 5 μg/ml doxycycline and 1 mM sodium butyrate. Cells were crosslinked in 2% formaldehyde for 10 min at room temperature, quenched in 0.125 M glycine for 5 min, and washed twice with ice-cold PBS. Crosslinked cell pellets were mixed with 1 mL ice-cold ChIP lysis buffer [50 mM HEPES pH 7.9, 140 mM NaCl, 1 mM EDTA, 10% glycerol, 0.5% NP-40, 0.25% Triton X-100, protease inhibitor (Roche)] and incubated rotating at 4°C for 10 min then spun at 1700 x g for 5 min at 4°C. Nuclei were resuspended in wash buffer [(10 mM Tris-HCl pH 7.5, 100 mM NaCl, 1 mM EDTA pH 8.0, protease inhibitor (Roche)] and rotated for 10 min at 4°C. Nuclei were collected by centrifugation at 1700 x g for 5 min at 4°C, then gently rinsed with shearing buffer (50 mM Tris-HCl pH 7.5, 10 mM EDTA, 0.1% SDS) followed by centrifugation at 1700 x g for 5 min at 4°C. After a second rinse with shearing buffer, nuclei were resuspended in 1 mL of shearing buffer and transferred to a milliTube with AFA fiber (Covaris). Chromatin was sheared using a Covaris S220 for 5 min (peak power: 140, duty cycle: 5, cycles/burst: 200).

Chromatin was spun at 16,000 x g for 10 min at 4°C and the pellet was discarded. The chromatin was pre-cleared with protein A + protein G beads blocked with 200 μg/mL glycogen, 200 μg/mL BSA, 200 μg/mL *E. coli* tRNA for 2 h at 4°C. Pre-cleared chromatin (25 μg) was diluted in shearing buffer to 500 μL, adjusted to include 150 mM NaCl and 1% Triton X-100, then incubated with 10 μg anti-HA antibody (Cell Signaling C29F4) or 10 μg rabbit IgG (Southern Biotech) overnight. Samples were rotated with 25 μL pre-blocked protein A + G beads (Thermo Fisher) for 2 h at 4°C. Beads were washed with low salt immune complex (20 mM Tris pH 8.0, 1% Triton X-100, 2 mM EDTA, 150 mM NaCl, 0.1% SDS), high salt immune complex (20 mM Tris pH 8.0, 1% Triton X-100, 2 mM EDTA, 500 mM NaCl, 0.1% SDS), lithium chloride immune complex (10 mM Tris pH 8.0, 0.25 M LiCl, 1% NP-40, 1% deoxycholic acid, 1 mM EDTA), and TE buffer (10 mM Tris-HCl pH 8.0, 1 mM EDTA) for 10 min each at 4°C with rotation. DNA was eluted from the beads using 100 μL of elution buffer (150 mM NaCl, 50 μg/ml Proteinase K) and incubated at 55°C for 2 h then at 65°C for 12 h. DNA was purified using an Oligo Clean & Concentrator kit (Zymo Research). Purified DNA was quantified by qPCR using iTaq Universal SYBR Mastermix (BioRad) and the indicated primers (Table 1) for 50 cycles. Each sample was normalized to its own input.

## ACKNOWLEDGEMENTS

We thank Divya Nandakumar for her helpful suggestions and critical reading of the manuscript. We thank Matthew Gardner for cloning the ORF66.stop and mutant rescue BACs. A.D. is The Rhee Family Fellow of the Damon Runyon Cancer Research Foundation (DRG-2349-18). B.G. is an investigator of the Howard Hughes Medical Institute. This research was also supported by NIH R01AI122528 to B.G.

## REFERENCES

1. Nandakumar D, Glaunsinger B. 2019. An integrative approach identifies direct targets of the late viral transcription complex and an expanded promoter recognition motif in Kaposi’s sarcoma-associated herpesvirus. PLoS Pathog 15:e1007774.

2. Tang S, Yamanegi K, Zheng ZM. 2004. Requirement of a 12-base-pair TATT-containing sequence and viral lytic DNA replication in activation of the Kaposi’s sarcoma-associated herpesvirus K8.1 late promoter. J Virol 78:2609–14.

3. Wong-Ho E, Wu TT, Davis ZH, Zhang B, Huang J, Gong H, Deng H, Liu F, Glaunsinger B, Sun R. 2014. Unconventional sequence requirement for viral late gene core promoters of murine gammaherpesvirus 68. J Virol 88:3411–22.

4. Aubry V, Mure F, Mariame B, Deschamps T, Wyrwicz LS, Manet E, Gruffat H. 2014. Epstein-Barr virus late gene transcription depends on the assembly of a virus-specific preinitiation complex. J Virol 88:12825–38.

5. Davis ZH, Hesser CR, Park J, Glaunsinger BA. 2016. Interaction between ORF24 and ORF34 in the Kaposi’s Sarcoma-Associated Herpesvirus Late Gene Transcription Factor Complex Is Essential for Viral Late Gene Expression. J Virol 90:599–604.

6. Davis ZH, Verschueren E, Jang GM, Kleffman K, Johnson JR, Park J, Von Dollen J, Maher MC, Johnson T, Newton W, Jager S, Shales M, Horner J, Hernandez RD, Krogan NJ, Glaunsinger BA. 2015. Global mapping of herpesvirus-host protein complexes reveals a transcription strategy for late genes. Mol Cell 57:349–60.

7. Gruffat H, Marchione R, Manet E. 2016. Herpesvirus Late Gene Expression: A Viral-Specific Pre-initiation Complex Is Key. Front Microbiol 7:869.

8. Brulois K, Wong LY, Lee HR, Sivadas P, Ensser A, Feng P, Gao SJ, Toth Z, Jung JU. 2015. Association of Kaposi’s Sarcoma-Associated Herpesvirus ORF31 with ORF34 and ORF24 Is Critical for Late Gene Expression. J Virol 89:6148–54.

9. Gong D, Wu NC, Xie Y, Feng J, Tong L, Brulois KF, Luan H, Du Y, Jung JU, Wang CY, Kang MK, Park NH, Sun R, Wu TT. 2014. Kaposi’s sarcoma-associated herpesvirus ORF18 and ORF30 are essential for late gene expression during lytic replication. J Virol 88:11369–82.

10. Nishimura M, Watanabe T, Yagi S, Yamanaka T, Fujimuro M. 2017. Kaposi’s sarcoma-associated herpesvirus ORF34 is essential for late gene expression and virus production. Sci Rep 7:329.

11. Castaneda AF, Glaunsinger BA. 2019. The Interaction between ORF18 and ORF30 Is Required for Late Gene Expression in Kaposi’s Sarcoma-Associated Herpesvirus. J Virol 93.

12. Pan D, Han T, Tang S, Xu W, Bao Q, Sun Y, Xuan B, Qian Z. 2018. Murine Cytomegalovirus Protein pM91 Interacts with pM79 and Is Critical for Viral Late Gene Expression. J Virol 92.

13. Wyrwicz LS, Rychlewski L. 2007. Identification of Herpes TATT-binding protein. Antiviral Res 75:167–72.

14. Brulois KF, Chang H, Lee AS, Ensser A, Wong LY, Toth Z, Lee SH, Lee HR, Myoung J, Ganem D, Oh TK, Kim JF, Gao SJ, Jung JU. 2012. Construction and manipulation of a new Kaposi’s sarcoma-associated herpesvirus bacterial artificial chromosome clone. J Virol 86:9708–20.

15. Martinez-Guzman D, Rickabaugh T, Wu TT, Brown H, Cole S, Song MJ, Tong L, Sun R. 2003. Transcription program of murine gammaherpesvirus 68. J Virol 77:10488–503.

16. Summers WC, Klein G. 1976. Inhibition of Epstein-Barr virus DNA synthesis and late gene expression by phosphonoacetic acid. J Virol 18:151–5.

17. Chakravorty A, Sugden B, Johannsen EC. 2019. An Epigenetic Journey: Epstein-Barr Virus Transcribes Chromatinized and Subsequently Unchromatinized Templates during Its Lytic Cycle. J Virol 93.

18. Notredame C, Higgins DG, Heringa J. 2000. T-Coffee: A novel method for fast and accurate multiple sequence alignment. J Mol Biol 302:205–17.

19. Laity JH, Lee BM, Wright PE. 2001. Zinc finger proteins: new insights into structural and functional diversity. Curr Opin Struct Biol 11:39–46.

20. Chen Y, Ciustea M, Ricciardi RP. 2005. Processivity factor of KSHV contains a nuclear localization signal and binding domains for transporting viral DNA polymerase into the nucleus. Virology 340:183–91.

21. Gruffat H, Kadjouf F, Mariame B, Manet E. 2012. The Epstein-Barr virus BcRF1 gene product is a TBP-like protein with an essential role in late gene expression. J Virol 86:6023–32.

22. Basehoar AD, Zanton SJ, Pugh BF. 2004. Identification and distinct regulation of yeast TATA box-containing genes. Cell 116:699–709.

23. Labo N, Miley W, Marshall V, Gillette W, Esposito D, Bess M, Turano A, Uldrick T, Polizzotto MN, Wyvill KM, Bagni R, Yarchoan R, Whitby D. 2014. Heterogeneity and breadth of host antibody response to KSHV infection demonstrated by systematic analysis of the KSHV proteome. PLoS Pathog 10:e1004046.

24. Gardner MR, Glaunsinger BA. 2018. Kaposi’s Sarcoma-Associated Herpesvirus ORF68 Is a DNA Binding Protein Required for Viral Genome Cleavage and Packaging. J Virol 92.

